# Engineering mutualism via nitrogen exchange in mixotrophic cocultures between *Clostridium acetobutylicum* and *Clostridium ljungdahlii*

**DOI:** 10.1101/2025.10.22.683918

**Authors:** Noah B. Willis, Paige A. Bastek, Aravind K. Arunachalam, Eleftherios T. Papoutsakis

## Abstract

We have previously shown that mixotrophic cocultures of *Clostridium acetobutylicum* and *Clostridium ljungdahlii* – using sugars and H_2_ as substrates – increase sugar-substrate carbon and electron conversion via CO_2_ and H_2_ capture and synthesize valuable products, such as isopropanol and 2,3-butanediol, that neither species can make independently. In this pairing, growth of *C. ljungdahlii* is constrained by *C. acetobutylicum*, since *C. ljungdahlii* relies on *C. acetobutylicum* to convert glucose into CO_2_, which *C. ljungdahlii* can use as a carbon and electron sources. However, this dependence is unilateral; *C. acetobutylicum’s* growth is not constrained by *C. ljungdahlii*. Consequently, population ratios between the two species can vary substantially throughout the course of fermentation and in different fermentation setups, typically with the faster growing *C. acetobutylicum* outcompeting *C. ljungdahlii*. Population ratio is an important variable because it influences metabolite yields and productivity and likely also impacts the initiation and frequency of the heterologous cell fusion events we have documented between *C. acetobutylicum* and *C. ljungdahlii*. Thus, developing methods to rationally control and maintain the population ratio are important for both biotechnological applications and fundamental study of this coculture pairing. In this study we show that the different nitrogen utilization capabilities of these two organisms enable engineering of a mutualistic mixotrophic syntrophy in which *C. ljungdahlii* relies on *C. acetobutylicum* for carbon and electrons and *C. acetobutylicum* relies on *C. ljungdahlii* for nitrogen. First, we confirm that *C. ljungdahlii*, but not *C. acetobutylicum*, can convert nitrate into biologically useful ammonium, enabling the design of a culture medium in which *C. acetobutylicum* can only grow in the presence of *C. ljungdahlii*. Second, we test different ratios of nitrate to ammonium in batch cocultures and demonstrate that rapid nitrate utilization by *C. ljungdahlii* prevents *C. acetobutylicum* from becoming nitrogen-limited at any point in batch fermentation. Finally, we show that feeding different rates of nitrate to cocultures in fed-batch mode enables control of the coculture growth rate, maintenance of stable population ratios, and higher isopropanol and butanol yields in cocultures between *C. acetobutylicum* and *C. ljungdahlii*.

## 1. INTRODUCTION

Natural syntrophic communities are stabilized by layers of mutualistic cross-feeding in which specialized organisms partner with one another to create highly efficient divisions of labor and maximize the unique capabilities of each species (1, 2). In contrast, the preferred approach in the era of recombinant DNA technology has been to genetically modify specific model microbes to produce nonnative products of interest in monocultures. This approach can work very well, especially for high value products that require minimal to moderate genetic engineering to produce; insulin bioproduction in *Escherichia coli* is a classic example (3). However, heterologous expression of large, complex biosynthetic pathways in a single microbe, which would be required to produce many products of interest, is often infeasible, even for model organisms. Many effective bioprocesses based on undefined, naturally occurring microbial consortia also exist, such as in wastewater treatment and food manufacturing, but these are limited to those which have been discovered in nature or have co-evolved with human society over time.

Theoretically, synthetic, defined cocultures present the best of both worlds by combining evolutionarily specialized organisms with little to no genetic modification in new-to-nature combinations tailored to maximize yield and production of target products. However, controlling the performance of these synthetic cocultures can be challenging. If one or more members of the consortia outcompete their partner(s), the synergistic benefits of the community are lost. Identifying and maintaining synergistic population balance (often in an unnatural context such as a bioreactor), is one of the most important and most difficult challenges to overcome in designing effective synthetic, defined microbial cocultures (4).

Here, we demonstrate how to leverage the different nitrogen utilization capabilities of two coculture partners, *Clostridium acetobutylicum* and *Clostridium ljungdahlii* to maintain stable population balance. Previously, we have shown that cocultures between *C. acetobutylicum* and *C. ljungdahlii* improve carbon recovery from sugar fermentation, can synthesize valuable products which neither species produces independently, and induce large interspecies exchanges of intercellular material, including protein, RNA, and DNA, via microbial cell fusion between the two coculture partners (5–7). In its original conception, this coculture is an example of “obligatory commensalism”: *C. ljungdahlii*, an acetogen, feeds on the hydrogen and carbon dioxide gases produced when *C. acetobutylicum* ferments glucose (5). Because *C. ljungdahlii* cannot consume glucose (8), *C. ljungdahlii* requires *C. acetobutylicum* to obtain the carbon it needs for growth, but *C. acetobutylicum* does not require *C. ljungdahlii* to grow.

To make the dependence mutual (“obligatory mutualism”), we introduced an additional layer of nitrogen cross-feeding. Both *C. acetobutylicum* and *C. ljungdahlii* can use ammonium as a nitrogen source. *C. ljungdahlii* can also grow by converting nitrate to ammonium, but *C. acetobutylicum* cannot metabolize nitrate. When fed more nitrate than it needs for its own growth requirements, *C. ljungdahlii* converts the excess nitrate into ammonium and secretes it into the growth medium (9) when the two species are cocultured, so that *C. acetobutylicum* can be fed a suitable nitrogen source. In this way *C. acetobutylicum* can be made to depend on *C. ljungdahlii* for nitrogen, just as *C. ljungdahlii* depends on *C. acetobutylicum* for carbon and electrons (Fig. 1A). In this study we describe the design and implementation of this mutualism-based control strategy and explore its impact on the growth rate, population ratio, and metabolic performance of cocultures between *C. acetobutylicum* and *C. ljungdahlii*.

**Fig. 1:**
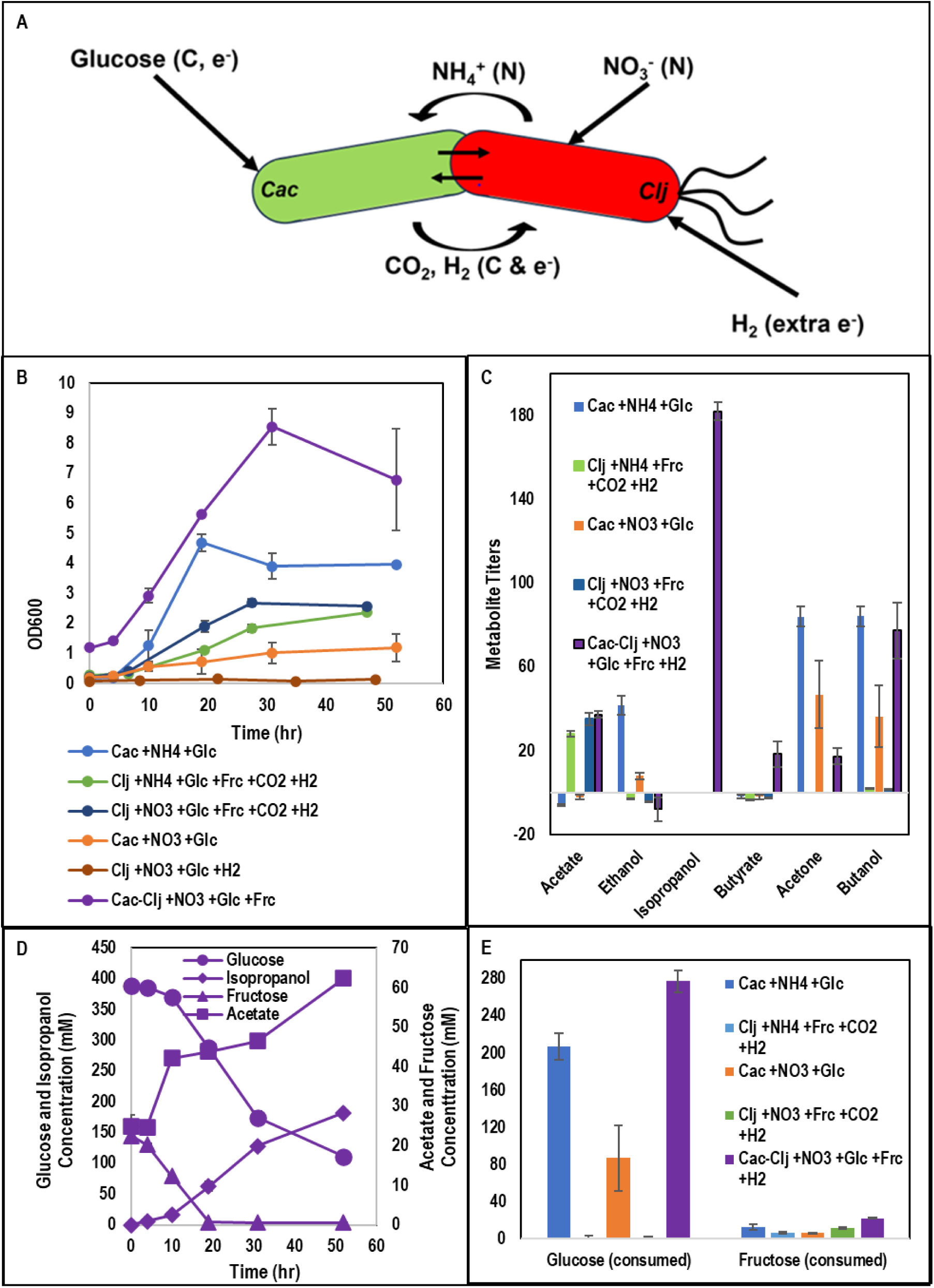
A) Schematic of engineered mutualism strategy. B) Growth kinetics for C. acetobutylicum and C. ljungdahlii monocultures grown in media with varied carbon, nitrogen, and electron sources and the C. acetobutylicum-C. ljungdahlii coculture grown with nitrate, glucose, fructose, and a H2 headspace. C) Metabolite titers for the monoculture and coculture varied carbon, nitrogen source experiments. D) Sugar consumption for the monoculture and coculture varied carbon and nitrogen, and electron source experiments. E) Glucose, fructose, acetate, and isopropanol kinetics for the C. acetobutylicum-C. ljungdahlii coculture grown with nitrate, glucose, fructose, and a H_2_ headspace. Error bars represent standard deviation between two biological replicates.

## 2. MATERIALS AND METHODS

### Plasmid and strain construction

Construction of all the *C. acetobutylicum* and *C. ljungdahlii* strains and associated plasmids used in this study have been previously described and are summarized in detail in Table S1.

### Microorganisms and growth media

*C. acetobutylicum* and *C. ljungdahlii* monocultures and cocultures were grown in either Turbo CGMB media (10) (for initial *C. acetobutylicum* tests on sodium nitrate in the presence of yeast extract and asparagine) or a newly designed defined, minimal synthetic growth medium designated “Minimal Turbo CGM with butyrate” or “TCGMinB.” TCGMinB was designed as a minimal, defined version of the previously developed Turbo CGM, a complex medium designed for *C. acetobutylicum-C. ljungdahlii* cocultures, and supplemented with butyrate due to butyrate’s growth benefit for *C. acetobutylicum* (11). On a per-liter basis, TCGMinB containes: 1.0 g KH_2_PO_4_, 1.25 g K_2_HPO_4_, 1.0 g NaCl, 0.01 g MnSO_4_·H_2_O, 0.348 MgSO_4_, 0.01 g FeSO_4_·7H_2_O, 0.20 Na_2_WO_4_·2H_2_O, 0.02 g CaCl_2_·2H_2_O, 8.0 mg of 4-aminobenzoic acid (PABA), 0.149 g L-methionine, 0.175 g cysteine-HCl·H_2_O, 2.46 g sodium acetate, 3.3 g sodium butyrate, 80 g glucose, and 3 g fructose. The medium was also supplemented with 10 mL each (per liter) of 100X trace element solution and 100x Wolfe’s vitamin solution (see medium DSMZ 879). For the inorganic nitrogen source, TCGMinB was supplemented with 10 mL per liter of either 200 g/L (NH_4_)_2_SO_4_ or 255 g/L NaNO_3_ to produce “TCGMinB-NH4” or “TCGMinB-NO3,” respectively, both of which contain 30 mM inorganic nitrogen from either ammonium or nitrate.

### Preparation of *C. acetobutylicum* and *C. ljungdahlii* for monoculture and coculture fermentations

For all *C. acetobutylicum* strains, a frozen stock was streaked onto a 2xYTG agar plate supplemented with 30mM of sodium butyrate and 40 μg/ml clarithromycin and left at 37°C in an anaerobic incubator for 3-5 days to allow for colony growth and spore formation. All subsequent pre-culturing steps were performed in TCGMinB-NH4 growth medium supplemented with 100 μg/ml clarithromycin. After plate incubation, a single colony was inoculated into a test tube containing 5 mL of growth medium (discussed below), heat shocked at 80°C to kill non-sporulated cells, supplemented with 100 μg/ml clarithromycin, and grown statically in an anaerobic incubator at 37°C for 16-24 hours. The resulting pre-culture was used to inoculate 30 mL of the same growth medium used for heat shock to an OD_600_ of 0.1. This culture grew statically in the anaerobic incubator at 37°C until it reached exponential phase (OD_600_ between 0.5-4.0). This pre-culture was used to inoculate the monoculture and/or co-culture.

For the initial test of *C. acetobutylicum’s* ability to grow on nitrate in the presence of yeast extract and asparagine, Turbo CGMB was used as the growth medium for pre-cultures. For all subsequent *C. acetobutylicum* monoculture and coculture experiments, TCGMinB-NH4 was used as the growth medium for *C. acetobutylicum* pre-cultures.

For *C. ljungdahlii* p100ptaHALO, a frozen stock was inoculated into 20 mL of YTF medium (12) supplemented with 3.5 g/L arginine and 25 g/L MES (“YTAF-MES”) (13) with 100 μg/ml clarithromycin in a 160 mL serum bottle which had been pre-flushed for 2 min with an 80% H_2_, 20% CO_2_ gas mix. The serum bottle was pressurized to 20 psig with the 80/20 gas mixture and grown for 24 hours at 37°C on a rotating platform at 90 rpm. This pre-culture was used to inoculate pre-flushed serum bottles each containing either TCGMinB-NH4 or TCGMinB-NO3 (depending on which inorganic nitrogen source would be used for the final culture) with 40 grams per liter of glucose and 100 μg/ml clarithromycin to an OD_600_ of 0.05-0.10. These pre-cultures were grown to exponential phase (OD_600_ of 0.6-1.2) and used to inoculate the monoculture and/or co-culture.

For *C. acetobutylicum-C. ljungdahlii* coculture preparation, volumes of exponential *C. acetobutylicum* and *C. ljungdahlii* pre-cultures corresponding to the desired target starting optical density and an R value of 5 (ratio of *C. ljungdahlii* to *C. acetobutylicum* cells based on OD_600_) (5) was centrifuged at 5000 rpm, 4°C for 10 min and then resuspended in 20 mL of Turbo CGMB in serum bottles pre-flushed with 100% H_2_. For batch serum bottle experiments, the target starting optical density was 1.0 OD_600_ and the initial serum bottle pressure was ∼10 psig. For fed-batch serum bottle experiments, the target starting optical density was 1.0 OD_600_ and the initial serum bottle pressure was ∼0 psig.

### Fed-batch sodium nitrate feeding to serum bottle cocultures

Fed-batch feeding of nitrate to serum bottle cocultures was accomplished using a 1600 Series Six Channel Syring Pump (New Era Pump Systems Inc., USA). Nitrate feed rates of 2.5, 1.3, and 0.63 mM per hour were achieved by delivering 0.46, 0.23, and 0.11M solutions of sodium nitrate at 0.11 ml per hour, respectively, to 20ml of culture volume in serum bottles. The sodium nitrate solutions were prepared by diluting a 3M sodium nitrate stock solution in TCGMinB-NO3 growth medium supplemented with 100 µg/ml clarithromycin.

### Metabolite and coculture population ratio analyses

High pressure liquid chromatography (HPLC) was used to quantify supernatant sugar and solvent concentrations as previously reported (14–16). For ammonium quantification, cell supernatants were diluted fivefold and analyzed on a YSI 2950D Biochemistry Analyzer using the installed ammonium biosensor module. Coculture population fraction of *C. ljungdahlii* p100ptaHALO was assayed via HaloTag (Promega, USA) labeling and flow cytometer (CytoFLEX S, Beckman Coulter Life Sciences, USA) analysis as previously described (17).

## 3. RESULTS & DISCUSSION

### 3.1. Development of a minimal, defined medium without organic nitrogen to test nitrate use by *C. acetobutylicum*

“Obligate mutualism” requires that neither organism in a partnership can grow without the presence of its partner. In coculture fermentations between *C. acetobutylicum* and *C. ljungdahlii* which use glucose as the sole carbon source, *C. ljungdahlii* absolutely depends on the CO_2_ generated by *C. acetobutylicum* for growth since, unlike *C. acetobutylicum*, *C. ljungdahlii* lacks the transporter required to grow on glucose (8). Previously, with respect to nitrogen, our coculture growth medium has included ammonium as the inorganic nitrogen source, a nitrogen form both *C. acetobutylicum* and *C. ljungdahlii* can use. *C. ljungdahlii* can also use nitrate for growth (via reduction to ammonium), assuming access to reducing equivalents from another substrate (such as H_2_, CO, or fructose). The genes which enable nitrate reduction to ammonium by *C. ljungdahlii* are well-characterized (9), and a simple BLAST search suggested that *C. acetobutylicum* does not possess this pathway. This observation inspired the design of an obligatory mutualism in which *C. ljungdahlii* depends on *C. acetobutylicum* to provide carbon (via oxidation of glucose to CO_2_) and *C. acetobutylicum* depends on *C. ljungdahlii* to provide nitrogen (via reduction of nitrate to ammonium) (Fig. 1A).

To confirm that *C. acetobutylicum* cannot grow on nitrate, we tested simple monocultures of *C. acetobutylicum* using a modified version of our typical growth medium (Turbo CGMB) in which the 15 mM of ammonium sulfate (30 mM total inorganic atomic nitrogen) was replaced with 30 mM of sodium nitrate (30 mM total inorganic atomic nitrogen). The *C. acetobutylicum* strain used for these experiments was CACas9 *hbd* p95ace02_atoB (10), a strain with the *hbd* gene knocked out (eliminating 4-carbon metabolism) and carrying a plasmid designed to maximize acetone production by overexpression of acetone pathway genes, genetic modifications unlikely to impact nitrogen metabolism.

Despite the replacement of ammonium sulfate with sodium nitrate, these *C. acetobutylicum* monocultures grew normally in the modified growth medium (Fig. S1). In hindsight, this was not surprising due to the significant quantity of organic nitrogen sources available in the medium. Though ammonium sulfate is the sole inorganic nitrogen source in our growth medium, the recipe also includes 5 g/L of yeast extract and 2 g/L asparagine. Though the precise composition of yeast extract can vary depending on the preparation process and manufacturer, estimates of the typical amino acid contents have been made with sufficient accuracy to enable a quantitative estimate of the total organic nitrogen content of our growth medium (18) (Supplementary Document 1). Based on this balance (which assumes the yeast extract composition in reference (18)), 15 mM ammonium sulfate (or sodium nitrate in the modified version) is responsible for only 36% of the total nitrogen in the growth medium; yeast extract (5 g/L) and asparagine (2 g/L) account for an additional 29% (24 mM) and 36% (30 mM), respectively. Since only 1.9 mM atomic nitrogen is required to build 1 OD_600_ of *C. acetobutylicum* biomass (Supplementary Document 1), our growth medium should be able to support a *C. acetobutylicum* cell density of up to approximately an OD_600_ of 27 using only the organic nitrogen sources (up to approximately 43 OD_600_ using both the organic and inorganic nitrogen sources). Since batch *C. acetobutylicum* monocultures typically only grow to an optical density of 8-14 in this growth medium (after which growth ceases due to solvent toxicity, pH inhibition, complete consumption of glucose, and/or transition to the stationary and sporulation growth phases), *C. acetobutylicum* growth never becomes limited by nitrogen in this medium, even in the absence of ammonium sulfate. This analysis suggested that testing the ability of *C. acetobutylicum* to grow on nitrate, and the proposed obligatory mutualism concept in coculture with *C. ljungdahlii*, requires the development of a minimal medium which uses nitrate as the sole nitrogen source.

Fortunately, defined growth media have previously been developed for both *C. acetobutylicum* (19) and *C. ljungdahlii* (20). Our Turbo Clostridial Growth Medium (“TCGMB”), which is designed to simultaneously support *C. acetobutylicum* and *C. ljungdahlii* (5, 10), already contains all components required for growth of *C. acetobutylicum* in the absence of supplemental amino acids. *C. ljungdahlii* has been shown to grow in a defined medium without yeast extract via supplementation of either 1 mM cysteine or 1 mM methionine (20). Therefore, to formulate a minimal medium which supports both organisms, we removed all of the yeast extract and asparagine from TCGMB and added 1mM each of cysteine and methionine, resulting in a minimal version of TCGMB which we dubbed “TCGMinB”. In TCGMinB the only nitrogen sources are the inorganic nitrogen source (ammonium sulfate or sodium nitrate) and 2 mM of total nitrogen from cysteine and methionine. Based on the aforementioned requirement for 1.9 mM nitrogen per OD600 of *C. acetobutylicum* biomass, the cysteine and methionine content of the medium can only support approximately 1.1 OD600 worth of *C. acetobutylicum* biomass, meaning that any *C. acetobutylicum* growth above this value must derive from inorganic nitrogen.

### 3.2. *C. ljungdahlii* enables *C. acetobutylicum* growth with nitrate as the sole nitrogen source

For the following experiments, we used the *C. ljungdahlii* DSM13528 strain transformed with the p100ptaHALO plasmid (21) which expresses the Halotag fluorescent reporter, enabling us to track *C. ljungdahlii* populations using flow cytometry. We initially tried to use the *C. acetobutylicum* ATCC 824 strain transformed with the p95_ZapA-FAST plasmid (22), which encodes an orthogonal fluorescent reporter and, in coculture with *C. ljungdahlii* carrying the Halotag, would have enabled direct monitoring of the *C. acetobutylicum* population and monitoring of interspecies cell fusion between *C. acetobutylicum* and *C. ljungdahlii* as we have previously demonstrated (6). However, for reasons that are unclear, the *C. acetobutylicum* strain transformed with p95_ZapA-FAST was unable to grow in minimal media, which was not the case with several other *C. acetobutylicum* strains tested (data not shown). Thus, we instead used *C. acetobutylicum* ATCC 824 transformed with the p95ace02a plasmid, a plasmid originally designed to maximize acetone production (11). By using p95ace02a in *C. acetobutylicum*, which shares the same antibiotic resistance marker as the p100ptaHALO used in *C. ljungdahlii* (MLS^R^), we could use clarithromycin to maintain the Halotag fluorescent reporter in *C. ljungdahlii* without killing *C. acetobutylicum*. This approach enabled direct monitoring of the *C. ljungdahlii* cell fraction and indirect determination of the *C. acetobutylicum* cell fraction (and thus the population ratio) by difference.

Using TCGMinB with ammonium sulfate (“TCGMinB-NH4”) as the nitrogen source, *C. acetobutylicum* grew quickly and to moderate cell density (up to 4.7 OD_600_) (Fig. 1B). However, when the nitrogen source was switched to sodium nitrate (“TCGMIN-NO3”), *C. acetobutylicum* growth was minimal and much slower, eventually reaching a peak cell density of 1.2 OD600 (Fig. 1B). As stated earlier, if monoculture of *C. acetobutylicum* cannot use nitrate, the maximum possible cell density in TCGMinB-NO3 should be 1.1 OD600 (from the small amounts of cysteine and methionine as previously discussed), a value which corresponds well with our experimental results if we allow for possible small errors in recording the optical density of *C. acetobutylicum* (which can form clumps of suspended biofilm) or carry over of small amounts of ammonium from precultures grown in TCGMinB-NH4. This result provided strong evidence that *C. acetobutylicum* cannot use nitrate as a nitrogen source for biomass formation. Notably, in coculture, *C. acetobutylicum* would be forced to compete with *C. ljungdahlii* even for the small amount of organic nitrogen available from cysteine and methionine, meaning that *C. acetobutylicum* growth on the small amount of organic nitrogen, when grown alongside *C. ljungdahlii,* is likely negligible in TCGMinB.

As expected, *C. ljungdahlii* grew well to typical cell densities (2-3 OD_600_) in both TCGMB-NH4 and TCGMB-NO3 with fructose (3 g/L) in the media and CO_2_ (as well as H_2_) in the headspace to serve as carbon and electron sources (Figure 1B). *C. ljungdahlii* actually grew slightly faster in the nitrate-containing version of TCGMinB, consistent with previous studies which have suggested that co-utilization of nitrate with hydrogen may increase the energy yield of hydrogen and enable faster acetogenic growth on CO_2_ (9, 23). However, when fructose was removed from the media and CO_2_ was removed from the headspace (glucose and H_2_ were still included), *C. ljungdahlii* could not grow in the nitrate media (Figure 1B), as expected, since it cannot use glucose as a carbon source.

Finally, we observed that cocultures of *C. acetobutylicum* and *C. ljungdahlii* grew quickly and to moderate-high cell densities (up to an OD_600_ of 8.6) in TCGMinB-NO3 (Fig. 1B), which uses nitrate as the sole inorganic nitrogen source and glucose as the major carbon source. Fructose (3 g/L) had to be included in this condition to enable *C. ljungdahlii* to begin to grow and metabolize nitrate, which, in turn, enabled *C. acetobutylicum* to grow and begin producing CO_2_ for *C. ljungdahlii*. However, all the fructose was exhausted by the 19-hour timepoint (Fig. 1D). After fructose depletion, the coculture continued to grow (2.9 OD_600_ worth of additional biomass), consumed an additional 177 mM of glucose (indicating strong ongoing *C. acetobutylicum* activity), and produced an additional 120 mM isopropanol and 18 mM additional acetate (both of which indicate strong ongoing *C. ljungdahlii* activity) (Fig. 1B, D), demonstrating obligate mutualism. Isopropanol (180 mM in total) was produced as the major product by the coculture, followed by butanol (79 mM) and acetate (37 mM) (Fig. 1C). Interestingly, more glucose was consumed by *C. acetobutylicum* in the coculture grown in TCGMinB-NO3 compared to the *C. acetobutylicum* monoculture grown in TCGMin-NH_4_ (Fig. 1E), even though both conditions were inoculated using the same *C. acetobutylicum* pre-cultures (we further explored and interpreted this finding in subsequent experiments).

### 3.3. Rapid nitrate conversion kinetics of *C. ljungdahlii* in batch cocultures prevent *C. acetobutylicum* from becoming nitrogen-limited

Our initial set of experiments successfully established that *C. acetobutylicum* monocultures cannot grow using nitrate as the nitrogen source. However, cocultures grown with nitrate as the sole nitrogen source (ignoring the negligible contribution of cysteine and methionine), grew well and consume glucose in large amounts, indicating that the presence of *C. ljungdahlii* enables *C. acetobutylicum* to use nitrate Presumably, *C. ljungdahlii* enables *C. acetobutylicum* growth by reducing excess nitrate to ammonium and passing it to *C. acetobutylicum* via secretion. Next, we designed an experiment to explore if we could tune the behavior and population ratio of the system by varying the nitrogen composition of TCGMinB. We hypothesized that, by growing the coculture in growth media with different nitrate:ammonium ratios, we could vary the extent to which *C. acetobutylicum* depends on *C. ljungdahlii* for nitrogen to grow, possibly modifying the growth rate, population ratio, and metabolic performance of the coculture.

To test this hypothesis, we performed cocultures in three different versions of TCGMinB. All three versions contained the normal starting amount of total inorganic nitrogen (30 mM), glucose (80 g/L), and fructose (3 g/L). However, the first version used ammonium sulfate as the sole nitrogen source, the second version used sodium nitrate as the sole nitrogen source, and the third version contained equal amounts of nitrogen from both nitrogen sources (Fig. 2A). Like the previous set, these experiments were conducted using *C. ljungdahlii* and *C. acetobutylicum* transformed with the p100ptaHALO and p95ace02a plasmids, respectively.

**Fig. 2:**
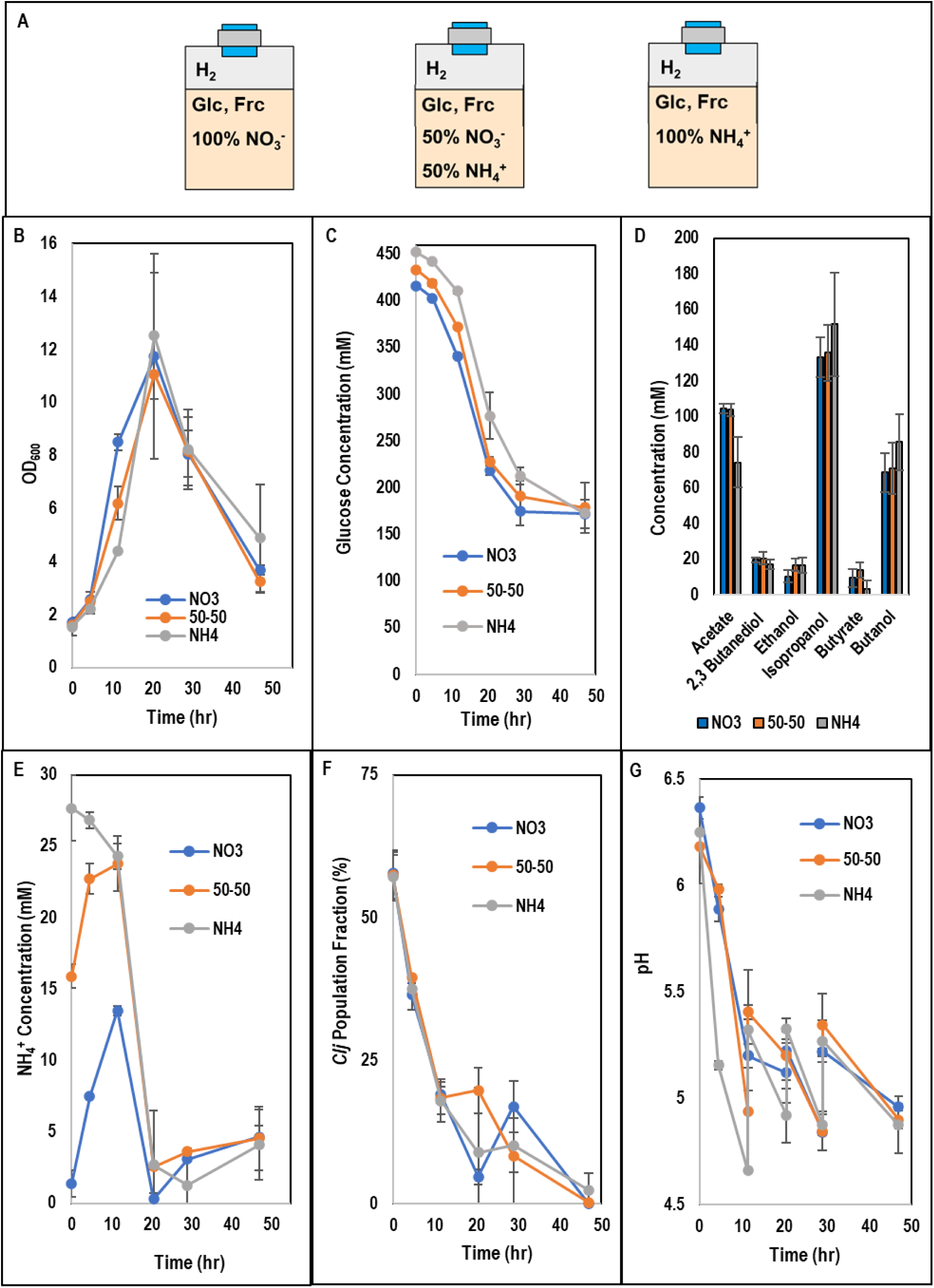
A) Schematic of growth medium composition for C. acetobutylicum-C. ljungdahlii grown in media with inorganic nitrogen composition of 100% NO3-, 50% NO3- and 50% NH4+, or 100% NH4+. B) Growth kinetics for the C. acetobutylicum-C. ljungdahlii cocultures grown with varied ammonium:nitrate ratios. C) Glucose consumption kinetics for the C. acetobutylicum-C. ljungdahlii cocultures grown with varied ammonium:nitrate ratios. D) Metabolite titers for the C. acetobutylicum-C. ljungdahlii cocultures grown with varied ammonium:nitrate ratios. E) Ammonium kinetics for the C. acetobutylicum-C. ljungdahlii cocultures grown with varied ammonium:nitrate ratios. F) C. ljungdahlii population fraction for the C. acetobutylicum-C. ljungdahlii cocultures grown with varied ammonium:nitrate ratios. G) pH profile for the C. acetobutylicum-C. ljungdahlii cocultures grown with varied ammonium:nitrate ratios. Error bars represent standard deviation between two biological replicates.

Contrary to our expectation, varying the nitrogen source ratio did not induce any statistically meaningful differences in peak biomass accumulation, glucose consumption, *C. ljungdahlii* population fraction, or metabolite production (Fig. 2B-D, F; Fig. S2) between the three different coculture conditions. Since the kinetics of *C. ljungdahlii* nitrate to ammonium conversion are relatively quick (9), we considered that the rate of conversion of nitrate to ammonium by *C. ljungdahlii* may have outstripped the rate of consumption of ammonium by *C. acetobutylicum* in the nitrate-containing conditions, especially early in the fermentation. This could lead to ammonium accumulation in both nitrate-containing conditions, and thus *C. acetobutylicum* would not practically experience any difference in nitrogen availability relative to the ammonium-only condition.

As hypothesized, assaying the fermentation samples for ammonium content showed ammonium accumulation for the first 12 hours in both nitrate-containing conditions (Fig. 1E). Even the coculture grown with 100% nitrate accumulated up to 13 mM ammonium (out of 30 mM total inorganic nitrogen) before the ammonium levels began to decrease after the 12-hour timepoint (Fig. 1E). Notably, *C. acetobutylicum* did experience ammonium limitation in the nitrate-only condition around the 21-hour timepoint. However, by this point in the fermentation, all the biomass accumulation and most of the glucose consumption had already occurred (Fig. 2B, C), so *C. acetobutylicum* was probably not significantly limited by the lack of ammonium. This result shows that *C. ljungdahlii* can rapidly convert nitrate to ammonium to supply *C. acetobutylicum* with nitrogen.

Interestingly, the coculture grown with only nitrate grew to a higher cell density (an OD_600_ of 8.5) over the first 12 hours than the coculture with a 50-50 mix (an OD_600_ of 6.2), and both nitrate-containing cocultures outgrew the ammonium-only cocultures (an OD_600_ of 6.2)in the first 12 hours (reaching an OD_600_ of 4.4), although the peak cell densities reached by all three cocultures were statistically indistinguishable (Fig. 2B). Since *C. acetobutylicum* grows faster than *C. ljungdahlii*, and *C. acetobutylicum* growth was unrestrained by nitrogen availability in the ammonium-only condition, we expected to observe either the opposite trend in early growth rates or (considering rapid conversion of nitrate to ammonium by *C. ljungdahlii*) no difference. Since the population ratios (assayed via flow cytometry using the Halotag fluorescent reporter in *C. ljungdahlii*) are essentially identical for the first 12 hours (comprising three timepoints) between the three conditions, the faster growth rate cannot be attributed to increased proliferation of just one species; both *C. acetobutylicum* and *C. ljungdahlii* grew faster over the first 12 hours in the nitrate-containing conditions, and both species grew fastest in the condition in which the most nitrate was supplied.

Nitrate supplementation, as mentioned earlier, has been shown to increase the growth rate of *C. ljungdahlii* on CO_2_ and H_2_ (9, 23). We did not, however, anticipate that receiving nitrogen from *C. ljungdahlii* would lead to an increase in *C. acetobutylicum’s* growth rate. We have previously reported transcriptomic and growth data suggesting that *C. acetobutylicum* growth likely benefits from in situ H_2_ removal by *C. ljungdahlii* (24). One possible explanation for the increase in *C. acetobutylicum* growth rate could be that chemotactic attraction of *C. acetobutylicum* to ammonia secreted by *C. ljungdahlii* causes *C. acetobutylicum* to associate more “tightly” with *C. ljungdahlii,* causing improved removal of H_2_ by *C. ljungdahlii* from *C. acetobutylicum’s* microenvironment.

Notably, the rate of decrease in the pH over the first 12 hours (prior to manual adjustment upwards with sodium hydroxide) was fastest in the ammonium-only condition and slowest in the nitrate-only condition (Fig. 2G). We observed this trend even though the rate of acetic acid production (the major acid product) is lowest in the ammonium-only condition (Fig. S2C). The elevated pH in the nitrate-containing conditions, despite higher acid production, is readily explained by increased production of ammonium (a strong base) from nitrate by *C. ljungdahlii* (Fig. 2E). This maintenance of higher pH levels may also explain (in part or in full) the difference in early growth rates between the three coculture conditions, as both *C. ljungdahlii* and *C. acetobutylicum* prefers a pH range from 5.0-6.0 (though both can grow at a pH below 5.0).

Although coculture performance was not meaningfully “tuned” by varying the nitrogen source ratio, these batch experiments demonstrated rapid nitrate-to-ammonium conversion by *C. ljungdahlii* which both supplied *C. acetobutylicum* with nitrogen and improved pH control of the fermentation. We hypothesized that, due to these rapid kinetics, coculture growth control (via nitrogen limitation of *C. acetobutylicum*) would require fed-batch feeding of nitrate at a precise rate such that it is immediately utilized, preventing ammonium accumulation. We explore this approach next.

### 3.4. Fed-batch nitrate drip feeding enables control of coculture growth rate and maintenance of stable population ratios between *C. acetobutylicum* and *C. ljungdahlii*

To enable controlled, fed-batch nitrate feeding, we used a syringe pump to feed a concentrated nitrate solution to serum bottle coculture fermentations. The syringe pump was placed on the top shelf of an upright incubator, and the serum bottles were placed on the rotary shaker below and connected to the syringe pump with vinyl tubing using luer lock connections (Fig. S3). Each serum bottle started with 21 mL of culture, and each serum bottle was fed the same volumetric feed rate (0.11 mL/hr) such that approximately 6 mL of volume was added to each bottle over the course of the experiment. Specific nitrate feed rates were achieved by modifying the nitrate concentration of this feed solution. Eight 1 mL samples (including the initial timepoint) were taken every 5-10 hours throughout the course of the fermentation, so the volume in each bottle was maintained approximately between 19-20 mL. Total sugar consumed and metabolites produced (after accounting for dilution) were normalized to the starting volume of 20 mL (after removal of the 1 mL sample at the 0-hr timepoint).

Three nitrate feed rates were chosen carefully to induce nitrogen limitation in *C. acetobutylicum*. The base TCGMinB-NO_3_ medium contains 30 mM total inorganic nitrogen which is enough to produce approximately an OD_600_ of 16 of *C. acetobutylicum* biomass. Considering that coculture fermentations typically produce 8-12 OD_600_ biomass in the first 12-18 hours, we estimated that a nitrate feed rate of 2.5 mM per hour (30 mM in 12 hours) would match or slightly exceed the rate of nitrogen utilization by the coculture during the exponential growth phase, after which nitrate and/or ammonium would likely accumulate, serving as a non-nitrogen limited control. The second nitrate feed rate was set to 50% of this baseline rate (1.3 mM/hr; 30 mM total nitrogen in 24 hours), and the third nitrate feed rate was set to 25% of this baseline rate (0.63 mM/hr; 30 mM total nitrogen in 48 hours). The second and third nitrate feed rates were designed to induce moderate and severe nitrogen limitation of *C. acetobutylicum* in the coculture, respectively. Hereafter, we will refer to the 2.5, 1.3, and 0.63 mM/hr feed rates as the “high,” “medium,” and “low” nitrate feed conditions for simplicity.

Applying these rationally chosen feed rates produced exactly the desired effect, demonstrating the potential of our obligate mutualism approach to control growth and maintain stable population dynamics in the coculture between *C. acetobutylicum* and *C. ljungdahlii*. Ammonium did not accumulate above 4 mM in any of the three culture conditions (except for a small bump up to 9-12 mM at 5 hours while the cells were experiencing a lag phase) until after 17 hours (Fig. 3D). Notably, these low levels were maintained in the medium feed condition until 25 hours and in the low feed condition until 32 hours (Fig. 3D).

**Fig. 3:**
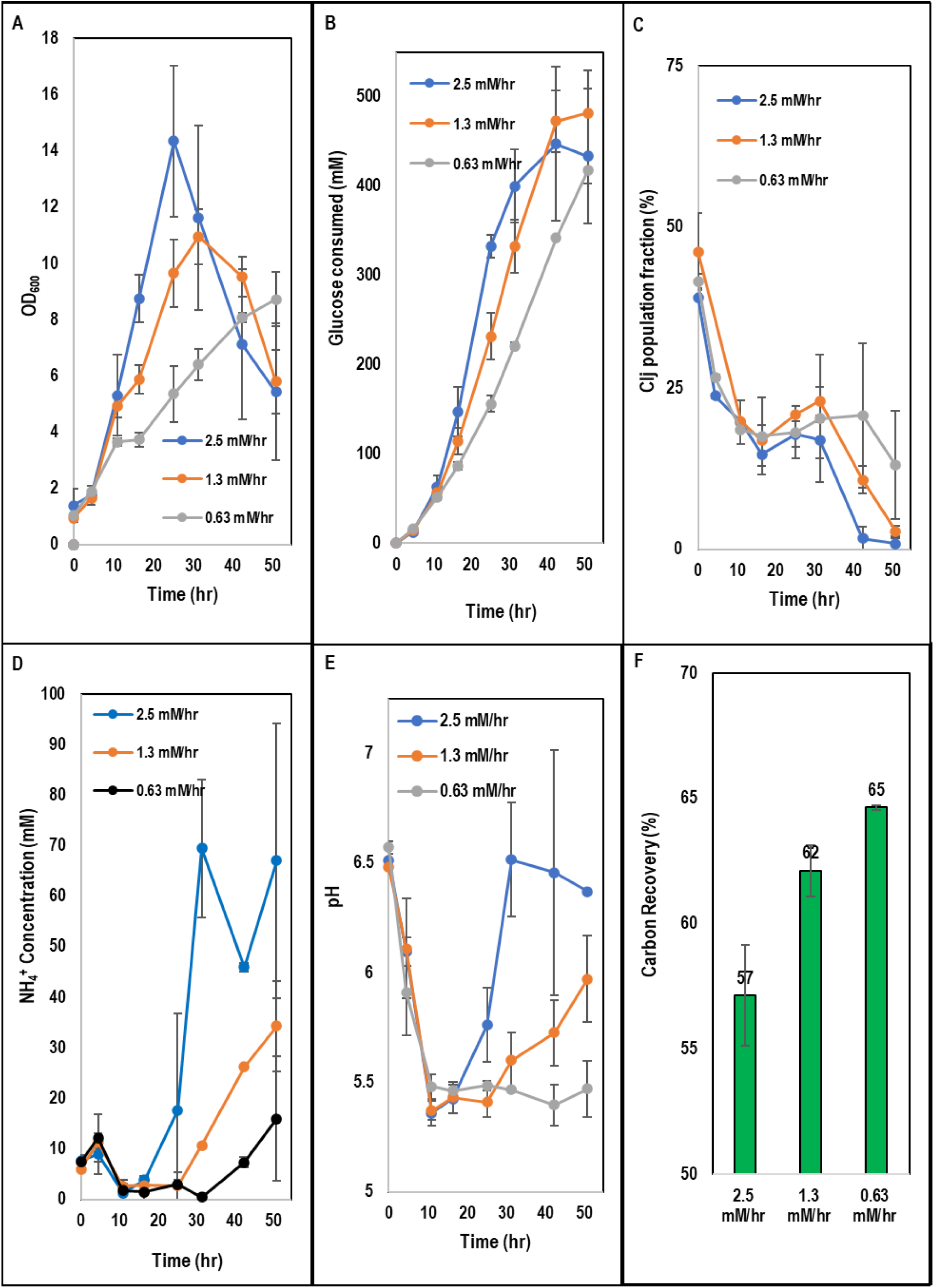
A) Growth kinetics from NO_3_^-^ fed-batch *C. acetobutylicum-C. ljungdahlii* cocultures. B) Glucose consumption kinetics from NO_3_^-^ fed-batch *C. acetobutylicum-C. ljungdahlii* cocultures. C) *C. ljungdahlii* population fraction kinetics from NO_3_^-^ fed-batch *C. acetobutylicum-C. ljungdahlii* cocultures. D) Ammonium kinetics from NO_3_^-^ fed-batch *C. acetobutylicum-C. ljungdahlii* cocultures. E) pH kinetics from NO_3_^-^ fed-batch *C. acetobutylicum-C. ljungdahlii* cocultures. F) Overall carbon recoveries from NO_3_^-^ fed-batch *C. acetobutylicum-C. ljungdahlii* cocultures. Error bars represent standard deviation between two biological replicates.

These ammonium profiles were clearly mirrored in the growth kinetics, glucose consumption rates, and pH behavior (Fig. 3A, B, E). Although early growth kinetics (up to 12 hours) were similar in all three conditions, the three conditions diverged after this point, showing fastest growth in the high nitrate feed condition and lowest growth in the low nitrate feed condition (Fig. 3A), exactly as expected. Biomass accumulation peaked at 25, 32, and 51 hours in the high, medium, and low feed rate conditions, respectively. In the case of the low feed rate condition, biomass is still accumulating at the end of the experiment (Fig. 3A). These growth dynamics are mirrored in the glucose consumption rates, where we observed fastest consumption in the high nitrate feed condition and slowest consumption in the low nitrate feed condition (Fig. 3B).

Regarding the pH, *C. acetobutylicum-C. ljungdahlii* batch fermentations typically require at least one manual pH adjustment to maintain pH levels above 5.0, where both species are most active (5). However, in this case, all three nitrate feed conditions maintained pH levels at or above 5.5 (Fig. 3E), likely due to in situ generation of ammonium from nitrate reduction by *C. ljungdahlii*. The medium and high nitrate feed rate conditions showed moderate, and large increases in pH after the initial drop to 5.5, corresponding to the accumulation of ammonium observed in those conditions (Fig. 3D, E).

The population ratios remained statistically indistinguishable between all three feed conditions until after 30 hours (Fig. 3C). Intuitively, one might have expected to observe an inverse correlation between the *C. ljungdahlii* population fraction and the nitrate feed rate since *C. acetobutylicum* growth is most restrained at lower nitrate feed rates. However, this thinking does not account for the fact that *C. ljungdahlii*, under the conditions tested here, is limited by the hydrogen production of *C. acetobutylicum*. To grow autotrophically, *C. ljungdahlii* requires H_2_ and CO_2_ in a 2-to-1 ratio (2 H_2_ per CO_2_). In this case, CO_2_ does not limit *C. ljungdahlii* since the cultures release excess, un-utilized CO_2_, as seen from the sub-100% carbon recoveries (Fig. 3F), and the bottle headspaces were flushed with pure H_2_ (and re-flushed after every sample) to maximize H_2_ availability. However, we have shown that, due to the low solubility of H_2_ and poor gaseous mass transfer in serum bottles mixed on rotary shakers (low k_L_a), relatively little of this headspace hydrogen is available in the liquid phase, causing *C. ljungdahlii* growth to be limited by H_2_ transfer in serum bottles (unpublished data). Thus, a large fraction of the H_2_ available to *C. ljungdahlii* in the coculture comes from glucose catabolism by *C. acetobutylicum* (H_2_ produced from glycolysis by *C. acetobutylicum* bypasses the headspace H_2_ mass transfer limitation because it is generated directly in the liquid phase). Therefore, when *C. acetobutylicum* growth and, critically, glucose consumption is limited by nitrogen availability, *C. ljungdahlii* also becomes limited by hydrogen availability from *C. acetobutylicum*, despite the fact that hydrogen is available in the headspace, because, as stated above, *C. ljungdahlii* growth is limited by low k_L_a and H_2_ solubility when grown in serum bottles. This bilateral limitation causes the coculture to form a stable population ratio, as we observe in this experiment, across multiple nitrate feed rates (Fig. 3C). This represents a strong demonstration of the utility of the obligate mutualism principle for establishing and maintaining a stable population ratio.

In terms of metabolite production and yield, nitrogen limitation of *C. acetobutylicum* favored carbon recovery and the production of alcohols, specifically isopropanol and butanol. The low nitrate feed condition increased the carbon recovery 14% relative to the high nitrate feed condition (Fig. 3F), likely due to the 9% and 32% higher production of 4-carbon metabolites (butyrate and butanol) and isopropanol, respectively (the medium feed condition demonstrated intermediate carbon recoveries and alcohol yields) (Fig. 4, Fig. S4). *C. acetobutylicum* 4-carbon metabolism enables ferredoxin reduction, increasing H_2_ production (10). Similarly, acetone production by *C. acetobutylicum* (which is converted to isopropanol by *C. ljungdahlii*) corresponds with higher H_2_ production (10). Since, as discussed earlier, *C. ljungdahlii* growth and CO_2_ fixation is limited by H_2_, increased H_2_ production by *C. acetobutylicum* (as indicated by increased butanol and isopropanol yields) enables increased CO_2_ fixation, producing higher carbon recovery in the conditions which produced the most isopropanol and butanol (Fig. 3F, Fig. 4, Fig. S4).

**Figure 4:**
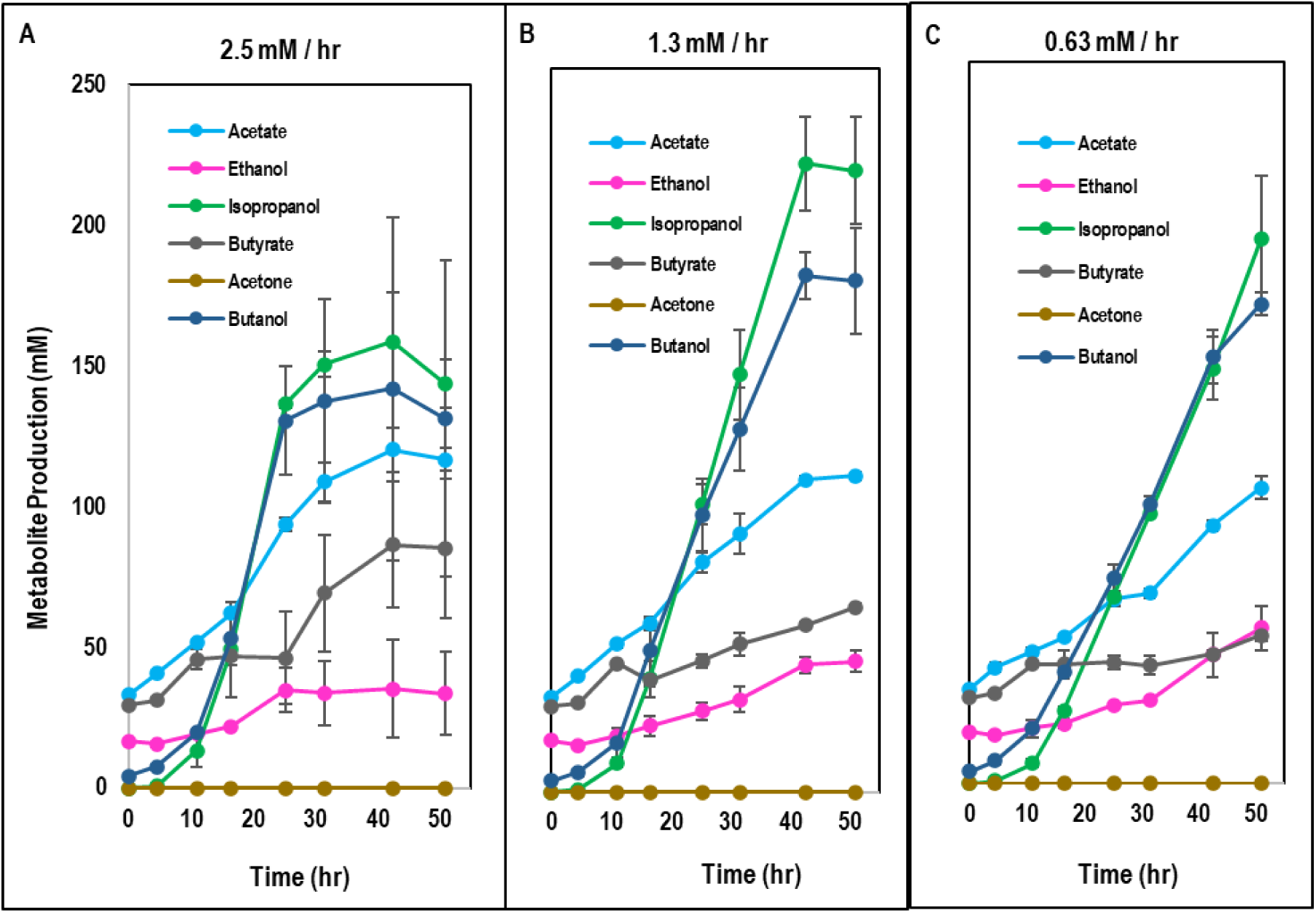
A) Metabolite kinetics for the C. acetobutylicum-C. ljungdahlii coculture fed NO3- at a rate of 2.5 mM/hr. B) Metabolite kinetics for the C. acetobutylicum-C. ljungdahlii coculture fed NO3- at a rate of 1.3 mM/hr. C) Metabolite kinetics for the C. acetobutylicum-C. ljungdahlii coculture fed NO3- at a rate of 0.63 mM/hr. Error bars represent standard deviation between two biological replicates.

The increased isopropanol (32%) and butanol (32%) yields from the low nitrate feed cocultures relative to the high nitrate feed cocultures (Fig. 4, Fig. S4) were likely due to one of two reasons (or a combination of both). Previous data from our group has suggested that ammonium limitation of *C. acetobutylicum* can produce increased yields of solvents such as acetone (converted to isopropanol by *C. ljungdahlii* in the coculture) and butanol (25). This is presumed to occur because, when *C. acetobutylicum* becomes nitrogen-limited, the cells cannot grow (or at least they cannot grow quickly) so they transition to stationary phase, and the *C. acetobutylicum* stationary phase favors solvent production. An alternative (or perhaps complementary) explanation is that chemotactic attraction to the ammonium produced by *C. ljungdahlii* causes *C. acetobutylicum* to associate more tightly with *C. ljungdahlii* cells in the cellular microenvironment. Our recent work suggests that increased cell-to-cell proximity to *C. ljungdahlii* induces increased per-glucose yields of acetone (isopropanol in coculture) by *C. acetobutylicum* (10). This effect was observed in a 4-carbon knockout *C. acetobutylicum* strain, so it does not speak to the increased butanol production, but it may help to explain the increased yield of isopropanol.

## 4. CONCLUSION

In this study, we show how a defined medium without organic nitrogen can be used to leverage the different nitrogen utilization capabilities of *C. acetobutylicum* and *C. ljungdahlii* and enforce obligate mutualism. Based on this approach, we demonstrate that controlled, drip-feeding of nitrate in fed-batch fermentation can control coculture growth, maintain stable population ratios, and increase carbon recovery and isopropanol and butanol yields. These results open several exciting avenues for further work, including rational manipulation of the coculture population ratio (by increasing reactor mixing such that *C. ljungdahlii* is no longer limited by *C. acetobutylicum* H_2_ production while *C. acetobutylicum* remains nitrogen-limited), use of a continuous reactor to maintain nitrogen limitation of *C. acetobutylicum* in coculture with *C. ljungdahlii* for prolonged periods of time, and measurement of heterologous cell fusion frequency under conditions in which both *C. acetobutylicum* and *C. ljungdahlii* must seek one another out to obtain essential nitrogen and carbon, respectively. These possibilities illustrate how engineering mutualism into synthetic microbial cocultures can expand our capacity to study interspecies microbial interactions and leverage the findings to solve real world problems.

## SUPPLEMENTAL MATERIAL

Supplementary Document 1 (Excel Sheet) (“Estimating Nitrogen Content from YE and ASN”)

Supplementary Figures (Word document) (Figures S1-S4)

## ACKNOWLEDGEMENTS

This work was supported by an ARPA-E project under contract AR0001505. N.B.W. was supported in part by a U.S. Department of Education GAANN Fellowship under grant P200A210065.

E.T.P. and N.B.W. conceived the project. E.T.P. and N.B.W. designed the experiments. N.B.W., P.A.B., and A. K. A. performed all experiments. E.T.P. and N.B.W. analyzed the data and wrote the manuscript.

---

**Table S1.**
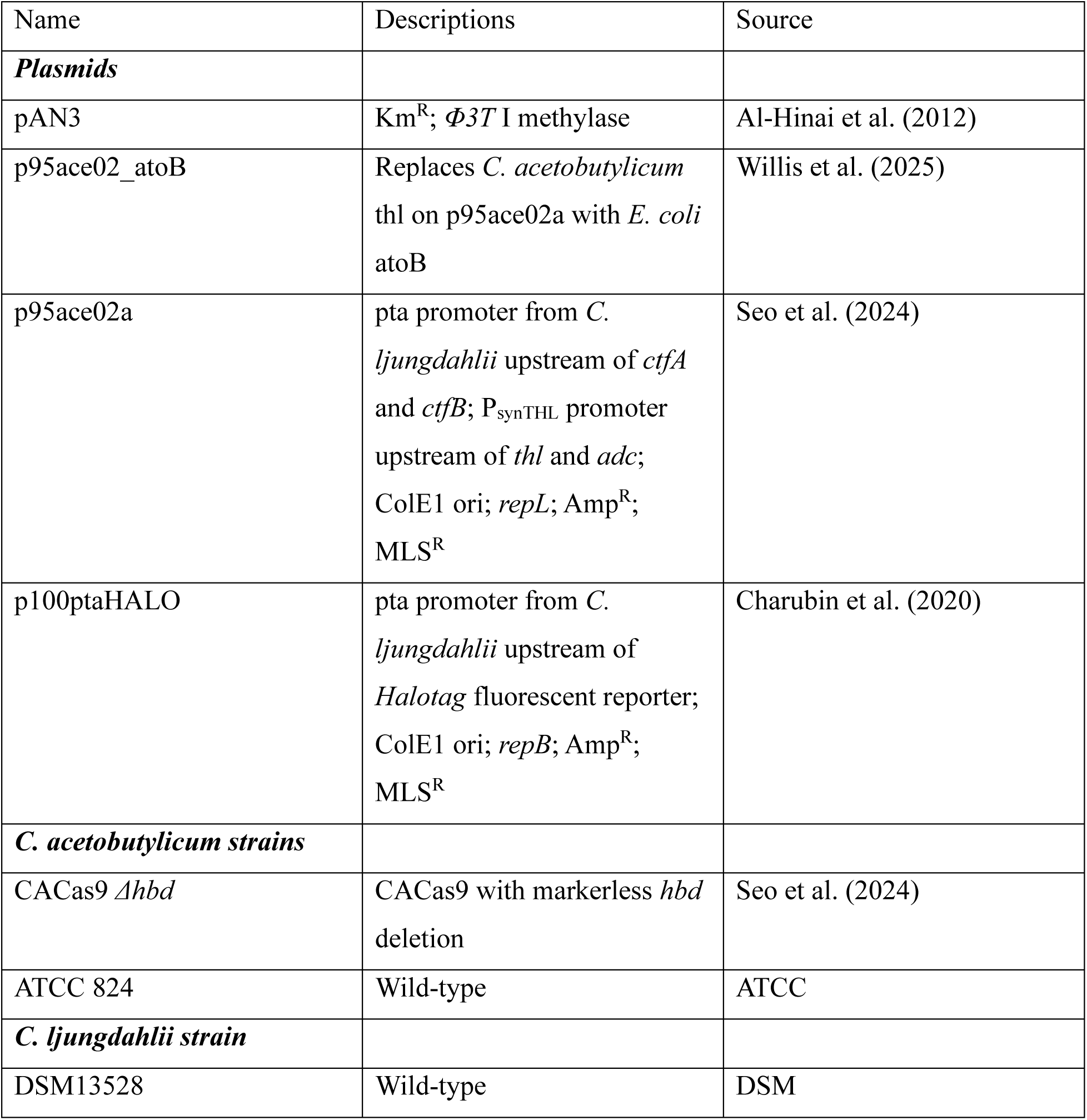
List of bacterial strains and plasmids used in this study.

| Name | Descriptions | Source |
| --- | --- | --- |
| ***Plasmids*** |  |  |
| pAN3 | Km<sup>R</sup>; $\Phi 3T$ I methylase | Al-Hinai et al. (2012) |
| p95ace02_atoB | Replaces *C. acetobutylicum* thl on p95ace02a with *E. coli* atoB | Willis et al. (2025) |
| p95ace02a | pta promoter from *C. ljungdahlii* upstream of *ctfA* and *ctfB*; P<sub>synTHL</sub> promoter upstream of *thl* and *adc*; ColE1 ori; *repL*; Amp<sup>R</sup>; MLS<sup>R</sup> | Seo et al. (2024) |
| p100ptaHALO | pta promoter from *C. ljungdahlii* upstream of *Halotag* fluorescent reporter; ColE1 ori; *repB*; Amp<sup>R</sup>; MLS<sup>R</sup> | Charubin et al. (2020) |
| ***C. acetobutylicum strains*** |  |  |
| CACas9 $\Delta hbd$ | CACas9 with markerless *hbd* deletion | Seo et al. (2024) |
| ATCC 824 | Wild-type | ATCC |
| ***C. ljungdahlii strain*** |  |  |
| DSM13528 | Wild-type | DSM |

**Fig. S1:**
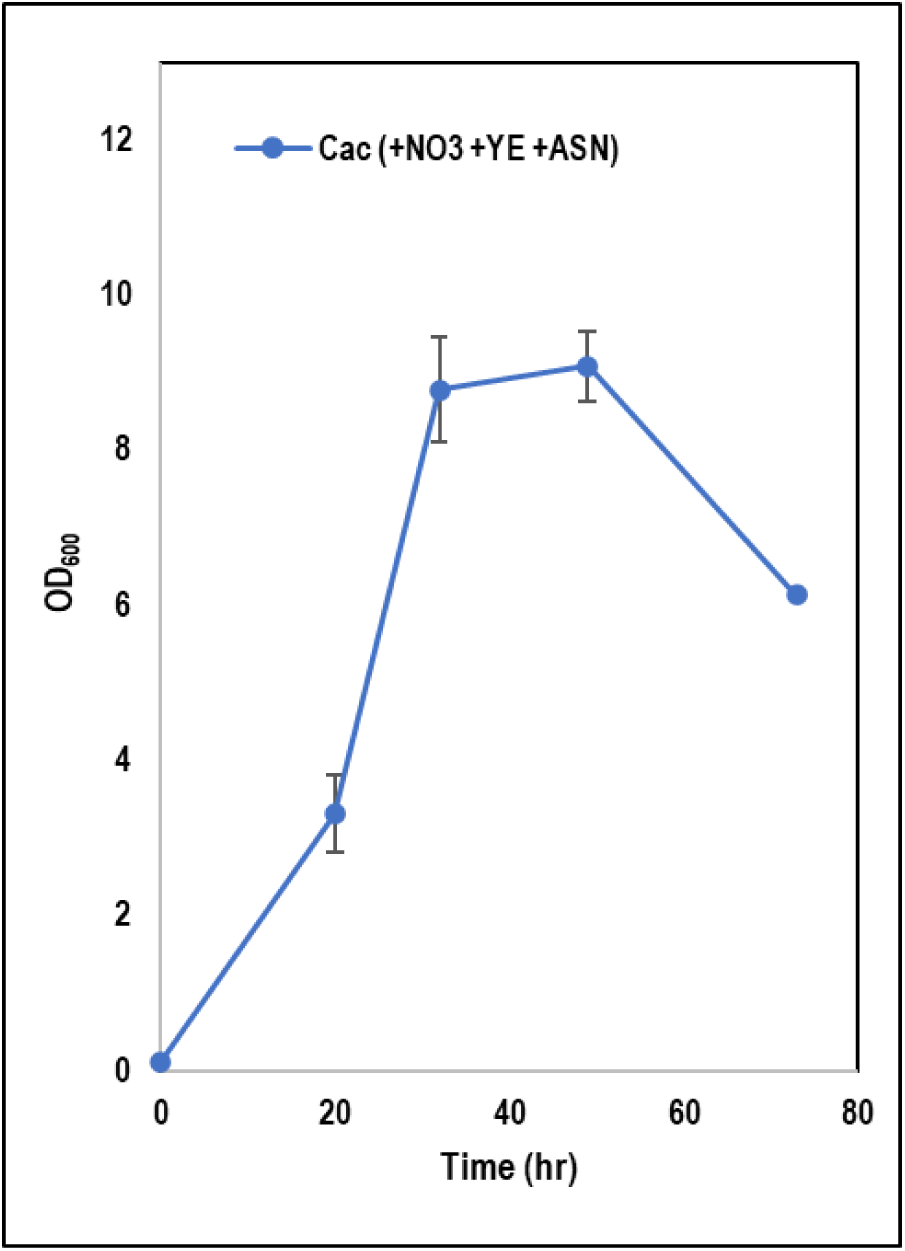
Growth kinetics for C. acetobutylicum grown in TCGMB medium with ammonium sulfate replaced with sodium nitrate. Error bars represent standard deviation between two biological replicates.

**Fig. S2:**
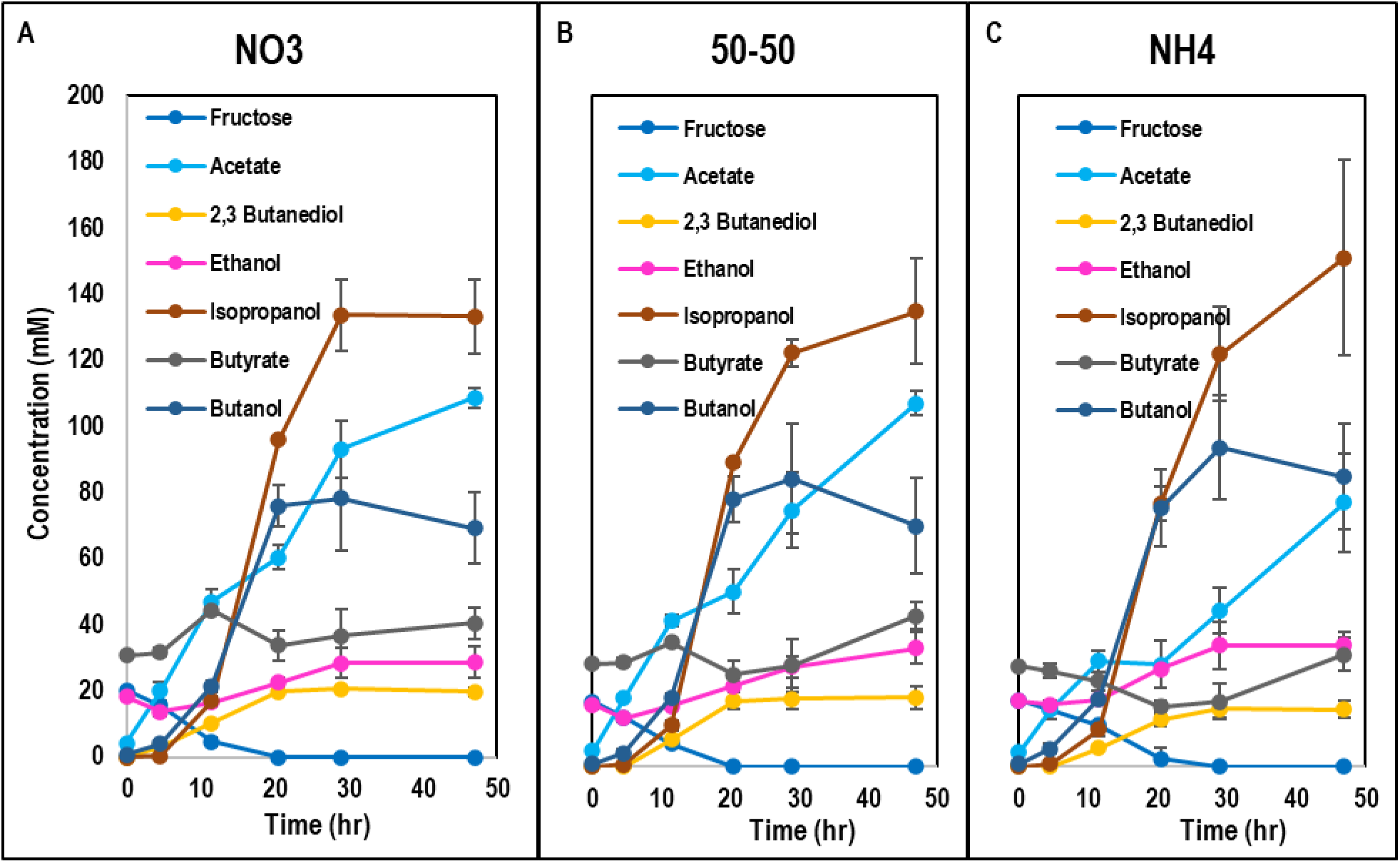
A) Metabolite kinetics for the *C. acetobutylicum-C. ljungdahlii* coculture grown in batch with 100% of inorganic nitrogen from NO_3_^-^. B) Metabolite kinetics for the *C. acetobutylicum-C. ljungdahlii* coculture grown in batch with 50% of inorganic nitrogen from NO_3_^-^ and 50% from NH_4_^+^. C) Metabolite kinetics for the *C. acetobutylicum-C. ljungdahlii* coculture grown in batch with 100% of inorganic nitrogen from NH_4_^+^. Growth kinetics for C. acetobutylicum grown in TCGMB medium with ammonium sulfate replaced with sodium nitrate. Error bars represent standard deviation between two biological replicates.

**Fig. S3:**
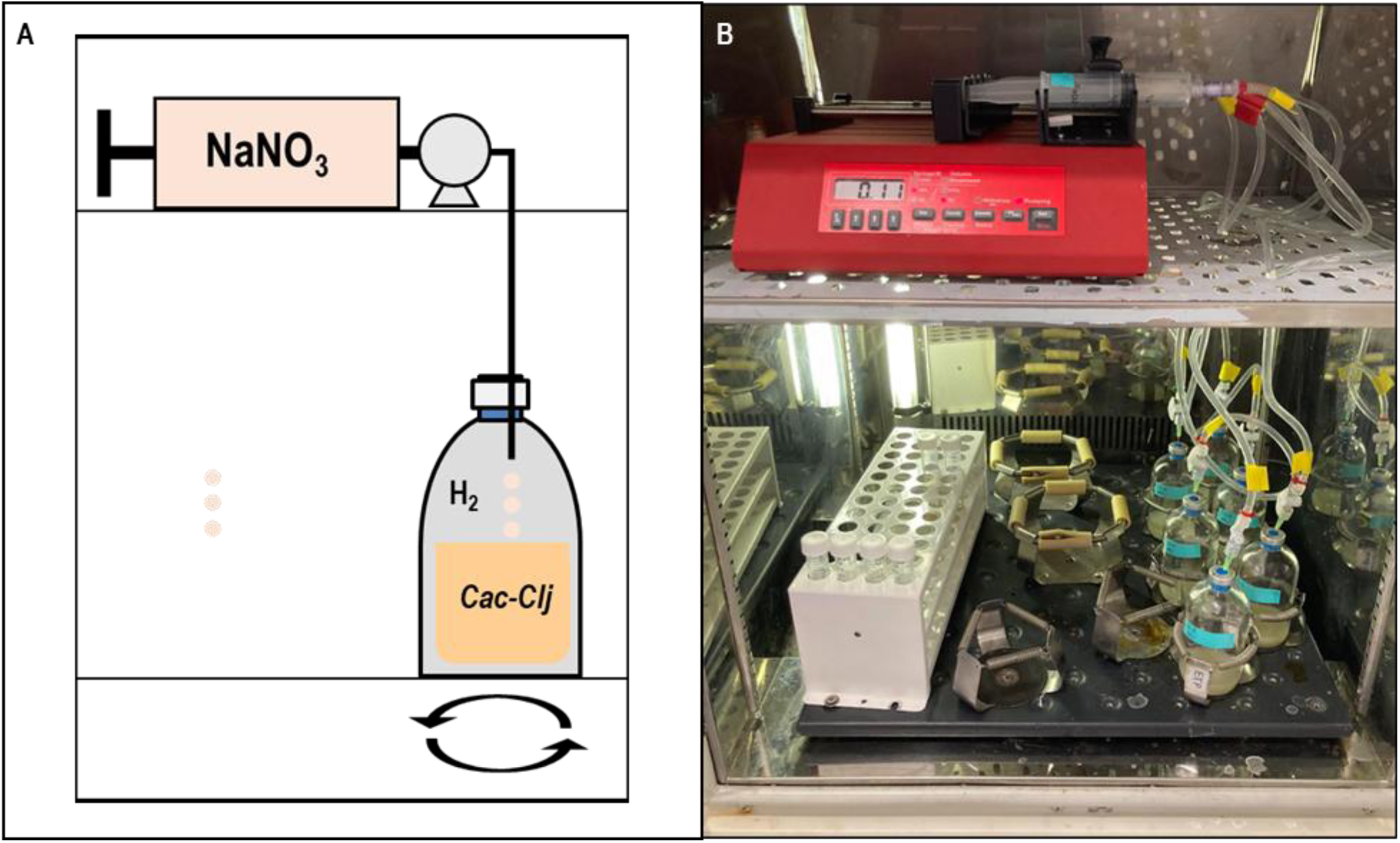
A) Schematic of NO_3_^-^ fed-batch experimental setup. B) Picture of NO_3_^-^ fed-batch experimental setup.

**Fig. S4:**
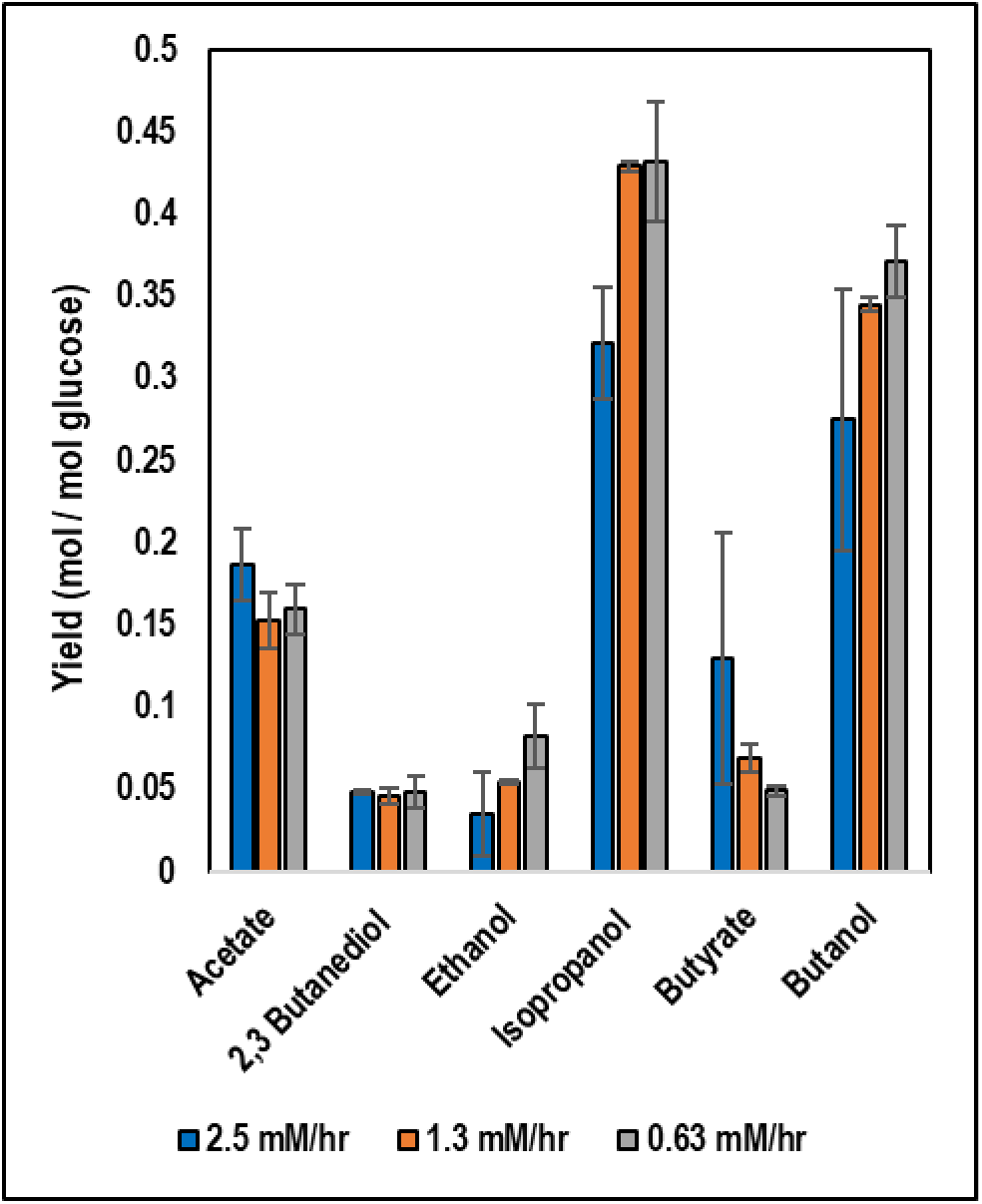
Metabolite yields for NO3-fed-batch C. acetobutylicum-C. ljungdahlii cocultures. Error bars represent standard deviation between two biological replicates.

## Notes

### Competing Interest Statement

The authors have declared no competing interest.

